# iRhom2 serves as a facilitator in obesity by enhancing adipose inflammation and insulin resistance

**DOI:** 10.1101/600460

**Authors:** Xu Minxuan, Ge Chenxu, Qin Yuting, Lou Deshuai, Li Qiang, Feng Jing, Wu Yekuan, Hu Linfeng, Huang Ping, Tan Jun

## Abstract

Chronic inflammation of adipose tissues contributes to obesity-triggered insulin resistance. Unfortunately, the potential molecular mechanisms regarding obesity associated systemic inflammation and metabolic disorder remain complicated. Here we display that inactive rhomboid-like protein 2 (iRhom2) is increased in mice fat with adipose inflammation. After 16 weeks on a high fat diet (HFD), obesity, chronic inflammation in adipose tissues and insulin resistance are markedly mitigated in iRhom2-knockout (iRhom2 KO) mice, but exaggerated in iRhom2-overactivated mice. The adverse impressions of iRhom2 on adipose inflammation and associated pathologies are determined in *db*/*db* mice. Also, we further exhibit that in response to HFD, iRhom2 KO mice and mice with deletion only in myeloid cells showed less severe adipose inflammation and insulin resistance than the control groups. Conversely, transplantation of bone marrow cells from normal mice to iRhom2 KO mice unleashed the severity of systemic inflammation and metabolic dysfunction after HFD ingestion. In conclusion, we identify iRhom2 as a key regulator that promotes obesity-associated metabolic disorder. Loss of iRhom2 from macrophages in adipose tissues inhibited the inflammation and insulin resistance. iRhom2 might be a therapeutic target for obesity-induced metabolic dysfunction.

**Significance:** Increased inactive rhomboid-like protein 2 signaling has recently been shown to trigger inflammation-associated activation of innate immune responses. Herein we investigate that this signal also plays a crucial role in obesity-triggered adipose tissue inflammation infiltration and metabolic disorder, beyond the well-known assignment in innate immune supervision. Also, we have reported the iRhom2 as a key promoter in regulating metabolic function, which enhances obesity-stimulated inflammation and systemic insulin resistance by up regulation of macrophages pro-inflammatory activation. Our current study indicates that targeting the iRhom2 signaling in adipose tissues could possibly be an efficient strategy to mitigating obesity-associated systemic inflammation and metabolic dysfunction.

## Introduction

Obesity, a major public health threat, which can cause metabolic diseases such as cardiac-cerebral vascular disease, cognitive dysfunction, type II diabetes and cancer is on the increase across the globe. Sterile inflammation induced by prolonged high fat diet intake in white adipose tissue (WAT), an important warehouse for energy-caching padding in the shape of triglyceride (TG), and for production of hormones and cytokines secretion in answer to homeostatic or nutritional alterations, exhibits a critical role in obesity-triggered systemic insulin resistance and its associated metabolic dysfunction (1-3). Unfortunately, the straightforward molecular mechanisms by which how nutritive obesity initiates systemic inflammation and metabolic disorder is still not fully understood.

In studies from the last few years, accumulating evidence has indicated an important role for innate immunity in the crosstalk with progression of obesity (4). As one of the most essential cells in adipose tissue associated innate immunity, adipose tissue macrophages have caught attention due to its core function in pro-inflammatory activation and connecting of metabolic dysfunction (5, 6). Nutritive obesity significantly promotes increased macrophage infiltration in adipose tissue, macrophages over-activation, as well as up regulated generation of pro-inflammatory cytokines and chemokines (such as tumor necrosis factor-α, TNF-α and monocyte chemoattractant protein-1, MCP-1) (7, 8). Thus, a series of regulators including nuclear factor κB (NF-κB), transforming growth factor activated kinase-1 (TAK1) and insulin receptor substract-1 (IRS-1) have been confirmed to participate in the inflammatory responses of macrophages and development of insulin resistance, which in turn crosstalk with adipocytes to protect against or promote adipose inflammation associated obesity. These foregone studies highlight the importance of innate immunity system, in particular of the role of macrophages in obesity (9-11). However, the great unknown is how these factors in innate immunity system markedly regulate metabolic balance, inflammation and its relationship in process of obesity identification.

Recently, inactive rhomboid protein 2 (iRhom2), also known as Rhbdf2, is an inactive member of the rhomboid intramembrane proteinase family of serine proteases, which have been shown to have crucial roles in regulating protein degradation, trafficking regulation and inflammatory response (12-14). With metabolic disorder or innate immunity changes, activity of iRhom2 in macrophages is significantly up regulated, and promotes TNF-α produce by trafficking of TNF-α converting enzyme (ADAM17), leading to activation of downstream pathway cascades involving tumor necrosis factor receptor 1/2 (TNFR1/2), TAK1, NF-κB and numerous cytokines or chemokines to accelerate inflammatory response and macrophages infiltration (15-17). These findings have indicated to several recent conclusions addressing how iRhom2 regulates immunoreaction stimulated by viral infection, arthritis and lupus nephritis (14, 18, 19). Of note, our preliminary study also indicated that iRhom2 exhibits an essential role in early kidney injury by regulation of inflammation and oxidative stress. Additionally, overexpression of iRhom2 as the promoter in the inflammatory responses contributes to acute hepatic injury and occurrence of systemic metabolism disorder (20). However, the correlation of iRhom2 to nutritive obesity associated adipose inflammation and insulin resistance has not been investigated. Also, dependent TNF-α activation and generation by iRhom2 only displays in immune cells (21). Given these findings, there is an imperious demand to explain the precise role for iRhom2 in the occurrence and development of obesity associated adipose inflammation and insulin resistance. The current work highlights and affords the primary demonstration to encourage the harmful role for iRhom2 in obesity, and this effects of iRhom2 is possibly performed by promoting macrophage activation of adipose and systemic inflammation.

## Results

### iRhom2 expression is increased in adipose tissues and involved in adipose inflammation and macrophages activation

To address the correlation of iRhom2 to adipose inflammation, we investigated iRhom2 activity in adipose tissues isolated from obese mice. The expression of iRhom2 were significantly up regulated in epididymal WAT (eWAT), inguinal WAT (iWAT) and purified adipocytes derived from iWAT or eWAT of high fat diet (HFD)-stimulated obese mice or *db/db* mice, which were associated with increased TNF-receptor 1 and 2. Consistent with this, in response to iRhom2 activation, obese mice displayed a higher activity of phosphorylation of NF-κB, as well as the pro-inflammatory cytokine of TNF-α in the downstream of iRhom2 than the same in SCD groups (Fig. 1 A-D and Fig. S1 A-C). Because adipose tissues contain not only adipose cells, but also macrophages and other type of cells. We further detected the alteration of iRhom2 expression in isolated F4/80 positive macrophages (MΦs) and stromal vascular fraction cells (SVFs) from adipose tissues. As expected, we found that HFD ingestion significantly elevated iRhom2 expression levels in MΦs derived from iWAT and eWAT, as well as the expression of iRhom2 downstream signaling activation including phosphorylation of NF-κB and TNF-α secretion (Fig. S1 D and E). Also, concurrent with the up regulation of iRhom2 activity in MΦs, obesity markedly elevated iRhom2/TNF-α/p-NF-κB signaling in SVFs collected from fat tissues (Fig. S1 F and G). These findings suggested that activation of iRhom2 and its downstream events in both macrophages and adipocytes may contribute to HFD-triggered inflammatory responses.

**Figure 1.**
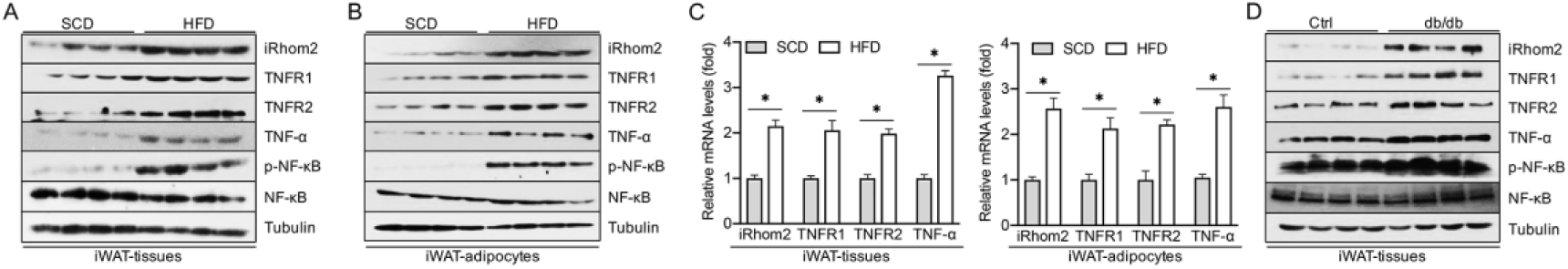
iRhom2 is increased in adipose tissues and promotes adipose inflammation in obese mice. (A) Western blotting analysis for expression of iRhom2, TNFR1/2, TNF-α and phosphorylated NF-κB in iWAT harvested from standard chow diet and high fat diet-fed C57BL/6N mice. (B) Western blotting analysis of isolated adipocytes in iWAT of SCD and HFD-fed C57BL/6N mice. (C) qPCR expression alteration for iRhom2, TNFR1/2 and TNF-α from iWAT or adipocytes in SCD and HFD-fed C57BL/6N mice; *n*=6. (D) Western blotting detection of iWAT collected from *db/db* mice and corresponding mice. Data were expressed as mean ± SEM, \**p* < 0.05.

### iRhom2 deletion protects against high fat diet-induced obesity and insulin resistance

Accumulating studies have indicated that over expression of iRhom2 and its downstream events may participate in the development of metabolic syndrome (17, 22). Herein we next examined the role of iRhom2 on HFD-induced metabolic alteration. The documentary body weight of mice was dramatically increased in both HFD-fed WT and iRhom2 knockout (KO) mice compared to SCD groups. Interestingly, after 16 weeks HFD feeding, iRhom2 KO mice, but not WT mice, exhibited a smaller body weight gain compared with WT groups (Fig. 2 A). Body composition examination also revealed a lower fat mass in HFD-fed iRhom2 KO mice than HFD-fed WT mice, although lean mass in HFD-fed iRhom2 KO mice did not differ markedly from those in HFD-WT mice (Fig. 2 B and C). Also, HFD-fed deficient mice did not display significant changes in food intake and water intake as did HFD-WT mice (Fig. S2 A-D). Of note, previous evidences have demonstrated that prolonged HFD feeding intake is able to promote insulin resistance and glucose intolerance (23-25). When obesity aspects were investigated, phenotypes of obesity and lipid accumulation in liver were significantly observed in HFD-fed WT mice compared with HFD-treated iRhom2 KO mice (Fig. 2 D), suggesting that iRhom2 deficiency may relieve HFD-induced obesity. In particular, time-course record (every 4 weeks) of blood glucose and insulin in HFD-fed iRhom2 KO mice were lower than those in WT mice, as well as the significant difference in area under curve (AUC) and HOMA-IR index (Fig. S2 E-G). Accordingly, in order to better explore this aspect, we further examined the insulin resistance signaling changes in HFD-fed KO mice and WT mice. As expected, following continuous HFD feeding for 16 weeks, the metabolism and insulin associated IRS-AKT-GSK3β and FOXO1-PEPCK pathway were investigated by western blotting, with alterations that the enhanced protein expression in phosphorylation of IRS1^Y608^, AKT, GSK3β, FOXO1 and increased PEPCK levels were significantly observed in the iWAT and eWAT samples of iRhom2 KO mice compared with the corresponding controls (Fig. 2 E). Consistently, iRhom2 KO mice displayed restored insulin sensitivity and improved glucose tolerance compared with WT mice as evidenced by glucose tolerance test (GTT) and insulin tolerance tests (ITT) detection (Fig. 2 F and G). Besides, what caught our attention was that HFD-fed iRhom2 KO mice exhibited up regulated fat mass and blood glucose, but displayed no significant difference in bone mineral density compared with those corresponding control mice (Fig. S2 H). Adipose or liver tissues were further subjected to hematoxylin and eosin (H&E) staining to visualize the impaired status of tissues (Fig. S2 I). iRhom2 KO had a slight difference on liver morphology in HFD-fed mice and markedly restrained a fatty liver morphology in mice compared with HFD-fed WT mice. Histological analysis also indicated that white adipocyte size and cells number were markedly reduced in the HFD-fed iRhom2 KO mice compared with WT mice (Fig. S2 J-L). In addition, immunohistochemical assay of F4/80 expression in adipose tissues revealed that iWAT and eWAT exhibited a lower F4/80 positive cells activity in iRhom2 KO mice than those in WT mice fed with HFD fodder, suggesting that iRhom2 expression in fat tissues may contribute to HFD-triggered macrophages activation and adipose inflammation infiltration (Fig. S2 M).

**Figure 2.**
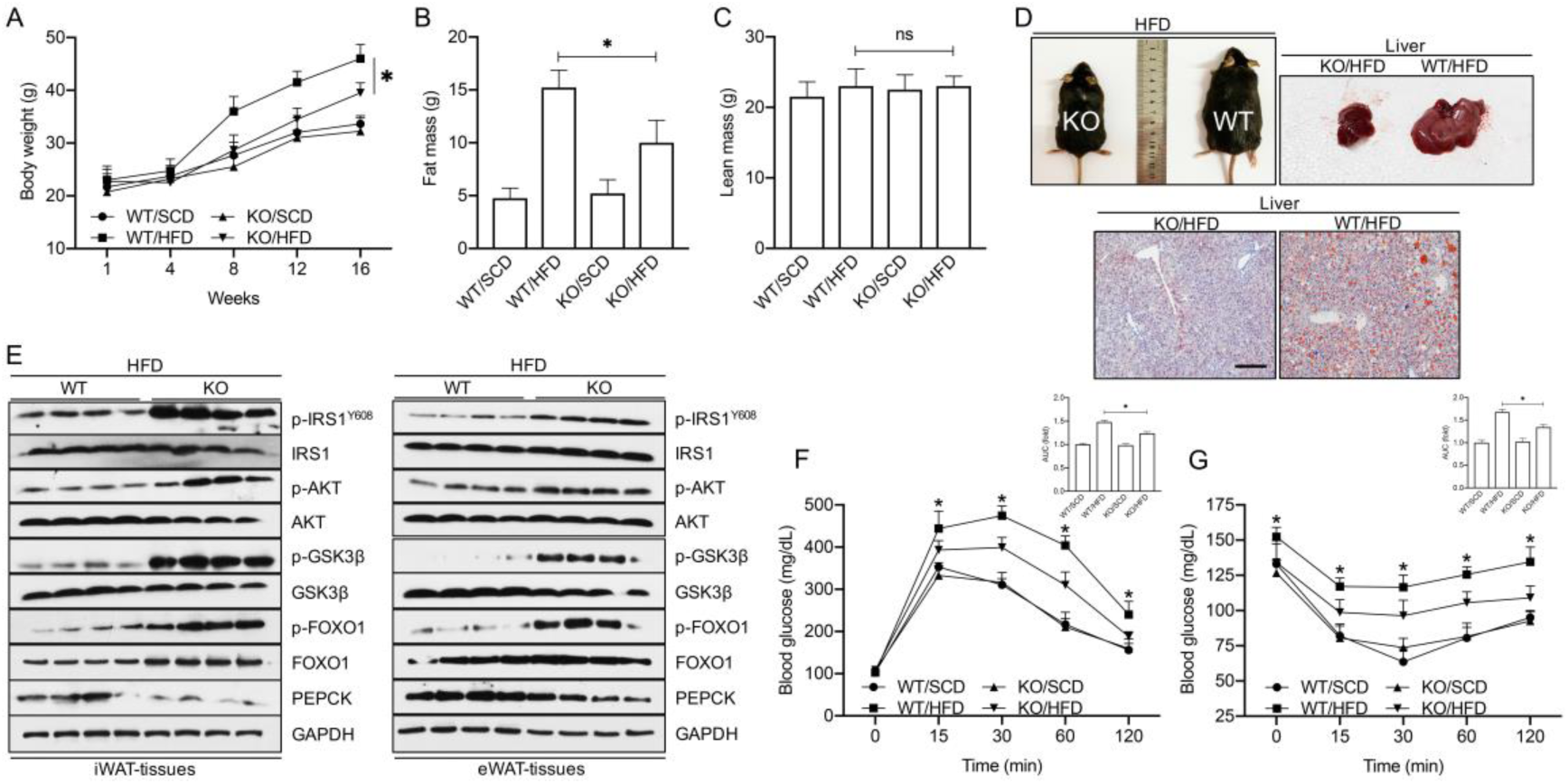
iRhom2 deficiency exhibits decreased fat mass and insulin resistance. (A) Body weight of male C57BL/6N WT mice and iRhom2 KO mice during the period of SCD and HFD feeding, *n*=15. (B) Fat mass and (C) lean mass examination was recorded in male C57BL/6N WT mice and iRhom2 KO mice with SCD and HFD feeding, *n*=15. (D) Representative experiment-recorded photos of HFD-fed WT mice, iRhom2 KO mice, appearance of the liver and oil-red O stained hepatic section (100 × magnification). (E) Representative immunoblot bands for levels of insulin resistance associated signaling expression including phosphorylated IRS1^Y608^, AKT, GSK3β, FOXO1 and PEPCK in iWAT and eWAT from HFD-fed WT and iRhom2 KO mice. (F) Glucose tolerance test and (G) insulin tolerance test of HFD or SCD-fed WT and iRhom2 KO mice, *n*=15. The data were expressed as the mean ± SEM. \**p* < 0.05 versus iRhom2 KO/HFD.

### Alteration of iRhom2 activity is coupled with high fat diet-triggered adipose inflammation and dyslipidemia

Given the close relationship between obesity, insulin resistance, inflammation and abnormal lipid metabolism, herein we next investigated the influence of iRhom2 in lipid metabolism and adipose inflammation. HFD-fed iRhom2 KO mice displayed a lower TNF-α/NF-κB signaling activation in iWAT compared with the same in HFD-fed WT mice (Fig. 3 A). Moreover, qPCR analysis of iWAT samples from mice undergoing HFD treatment revealed that the mRNA expression levels of genes associated with cholesterol synthesis (sterol regulatory element-binding protein 1c, SREBP-1c and 3-hydroxy-3-methyl-glutaryl-coenzyme A reductase, HMGCR), fatty acid uptake (fatty acid transport protein 1, FATP1; fatty acid binding protein, FABP1 and CD36) and fatty acid synthesis (fatty acid synthase, FAS; stearoyl-CoA desaturase, SCD1; acetyl CoA carboxylase α, ACCα and peroxisome proliferator-activated receptor γ, PPARγ) were markedly inhibited, whereas mRNA expression levels of genes involved in cholesterol efflux (ATP-binding cassette sub-family G member 1, ABCG1 and cholesterol 7α-hydroxylase, CYP7A1) and fatty acid β-oxidation (pyruvate dehydrogenase kinase 4, PDK4; peroxisome proliferator-activated receptor α, PPARα; carnitine palmitoyltransferase 1α, CPT-1α; acyl-coenzyme A oxidase 1, ACOX-1; long-chain acyl-CoA dehydrogenases, LCAD; medium-chain acyl-CoA dehydrogenase, MCAD and uncoupling protein 2, UCP2) were dramatically up regulated in iRhom2 KO mice relative to corresponding WT mice (Fig. 3 B). Consistently, reduced activation of inflammation associated mRNA levels in iWAT of HFD-fed iRhom2 KO mice including TNF-α, IL-1β, IL-6, MCP-1, iNOS, CXCR1, CCL1, IL-8, FKN, MDC, MIP-1, COX-2, CX3CR1, IL-18, CXCR4, ICAM1, Col4, TNFR1/R2, CCR2 and CXCL2 (Fig. 3 C), as well as down regulated serum levels of inflammatory chemokines and cytokines (Fig. S3 A) such as TNF-α, IL-1β, IL-6, MCP-1, CCL1, IL-18, IL-17, IL-2, IL-10, IL-5, MIP-2, MIP-3β, MCP-3, PCT, HMGB1, MDC, GCP-2, CX3CL1, CXCL10 and CXCL12 were significantly observed compared with those in HFD-fed WT mice. Of note, a lower TNF-α/NF-κB signaling activation in adipocytes and MΦs isolated from iWAT was also determined as evidenced by western blotting analysis (Fig. 3 D and E). Similarly, indicators’ examination (Fig. S4 A-E) in eWAT displayed a nearly consistent pattern of expression of those inflammation and lipid metabolic indicators relative to iWAT in HFD-fed iRhom2 mice, as indicated by qPCR and western blotting assay. The above results demonstrated that iRhom2 may directly contribute to HFD-induced adipose inflammation and dyslipidemia by partly, enhancement of adipocyte inflammation and macrophages activation. In order to better understand this aspect, mice were treated with adeno associated virus (AAV) packaged iRhom2 vectors to generate iRhom2 over-activated mice. As expected, after long-term HFD intake, an excess activation of iRhom2 in mice promotes to adipose inflammation, dyslipidemia development (Fig 3 F-J) and increased serum inflammatory cytokines and chemokines produce (Fig. S3 B). Also, the alteration of those indicators levels in eWAT were next examined as evidenced by qPCR and western blotting analysis. Collectively, these results significantly suggest that iRhom2 is a key linker involved in HFD-stimulated adipose inflammation and dyslipidemia.

**Figure 3.**
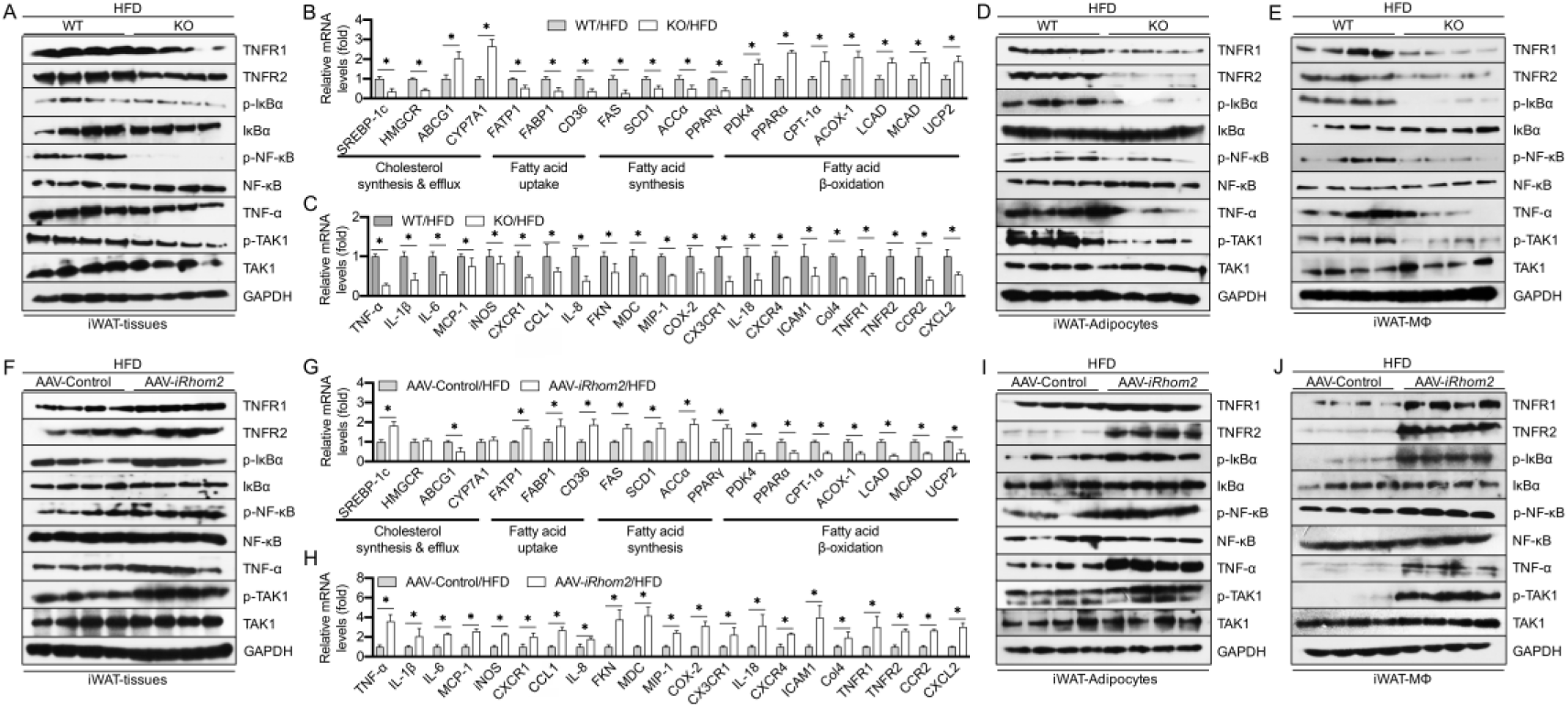
iRhom2 contributes to adipose inflammatory responses in of obese mice. (A) Representative immunoblot bands for expression of inflammation related signaling including TNFR1/2, phosphorylated IκBα, NF-κB, TAK1 and TNF-α. qPCR analysis detection for mRNA expression of genes responsible for (B) lipid metabolisms and (C) inflammatory cytokines and chemokines in HFD-fed WT mice and iRhom2 mice, *n*=6. Representative western blotting bands for expression of inflammatory signaling in (D) adipocytes and (E) F4/80+ macrophages isolated from iWAT of HFD-fed WT mice and iRhom2 mice. (F) Western blotting analysis for inflammatory signaling in iWAT of HFD-fed AAV-iRhom2 over-activated mice and corresponding mice. mRNA expression of genes responsible for (G) lipid metabolisms and (H) inflammatory cytokines and chemokines in HFD-fed AAV-control mice and AAV-iRhom2 over-activated mice, *n*=6. Western blotting for expression of inflammation indicators in (D) adipocytes and (E) F4/80+ macrophages from iWAT of AAV-control mice and AAV-iRhom2 over-activated mice. Data were expressed as mean ± SEM, \**p* < 0.05.

### Myeloid cell-specific iRhom2 knockout reduces high fat diet-induced adipose inflammation and dyslipidemia

The above data have extruded the role of iRhom2 in HFD-induced adipose inflammation and macrophages activation. In view of such situation, we attempted to deeply explore the role for iRhom2 in macrophages in adipose inflammation using bone marrow transplantation (BMT), a method that has been widely used for changing gene expression of macrophages (26, 27). The lethally irradiated WT mice were transplanted with bone marrow cells isolated from iRhom2 KO mice and/or WT mice to generate a KOBM→WT mice. After treatment of HFD for 16 weeks, the KOBM→WT/HFD mice, in which iRhom2 was deleted only in myeloid cells, and WTBM→WT/HFD mice, in which iRhom2 was integrated in all cells, obtained similar body weight and displayed a similar body composition (Fig. 4 A and B). Interestingly, KOBM→WT/HFD mice exhibited improved insulin sensitivity and restored glucose tolerance compared with WTBM→WT/HFD mice as evidenced by glucose tolerance test (GTT) and insulin tolerance tests (ITT) exanimation, and AUC index (Fig. 4 C and D). Accordingly, when insulin resistance associated indicators expression of iWAT were examined, KOBM→WT/HFD mice showed reduced severity of insulin resistance signaling including p-IRS^Y608^, p-AKT, p-GSK3β, p-FOXO1 and PEPCK levels compared with those in WTBM→WT/HFD mice (Fig. 4 E). When macrophages activation of iWAT were detected, indeed, KOBM→WT/HFD mice also displayed reduced TNF-α/NF-κB signaling activation, as indicated by western blotting (Fig. 4 F). Furthermore, indicated by changes in mRNA expression levels regarding lipid metabolism and adipose inflammation, KOBM→WT/HFD mice exhibited down regulated severity of dyslipidemia related indicators compared with WTBM→WT/HFD mice (Fig. 4 G). Also, WTBM→WT/HFD mice expressed dramatically more inflammation cytokines and chemokines in mRNA levels compared with those of KOBM→WT/HFD mice (Fig. 4 H). These results indicate that iRhom2 deficiency specifically in myeloid cells (MΦs) reduces the severity of HFD-induced obesity and insulin resistance.

**Figure 4.**
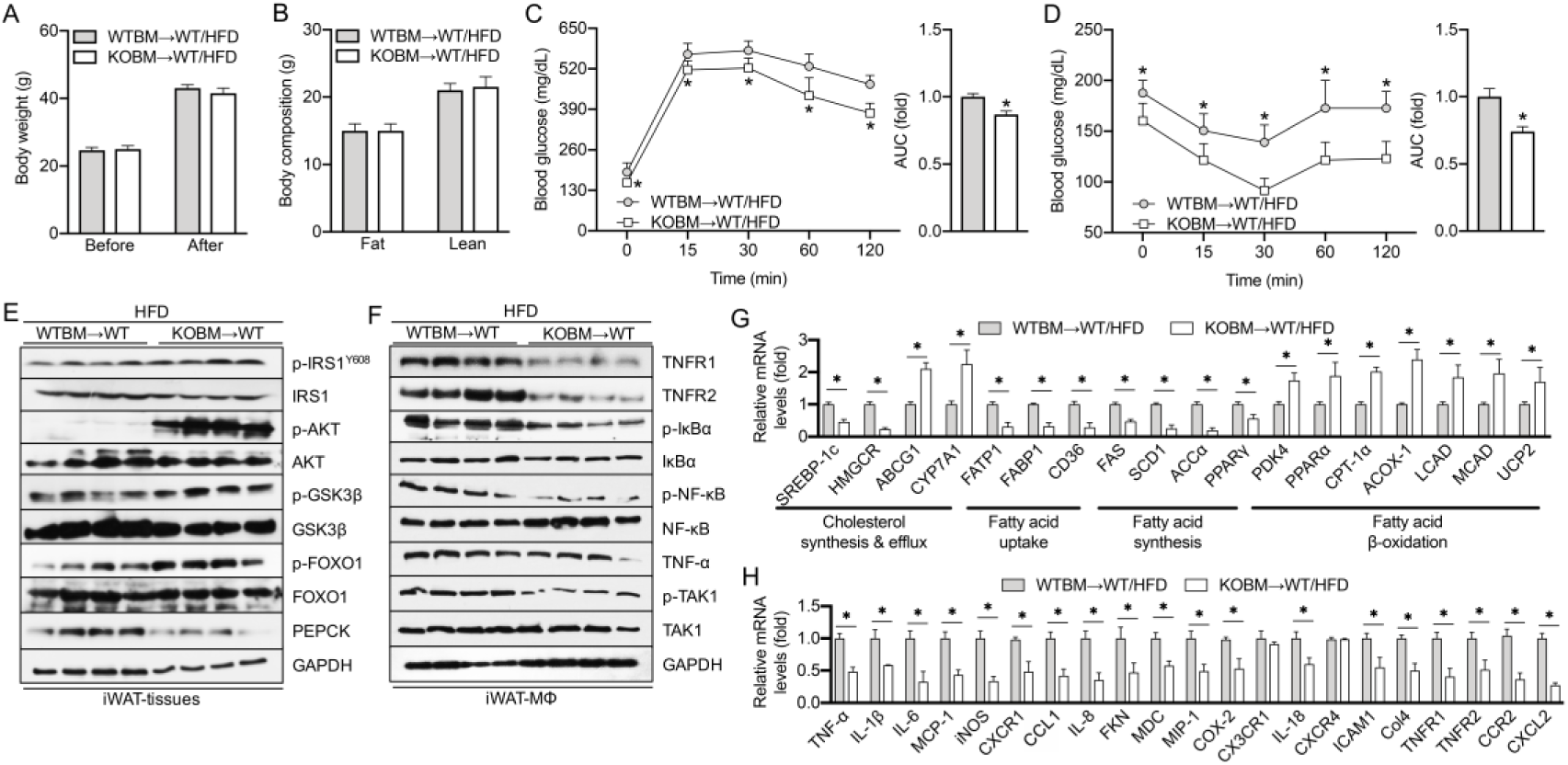
Myeloid cell-specific iRhom2 deletion down regulates insulin resistance and inflammation in obese mice. (A) Body weight was examined before and after the feeding period, *n*=15. (B) Body composition, *n*=10. (C) Glucose and (D) insulin tolerance test, *n*=15. Western blotting bands for expression of (E) insulin signaling from iWAT and (F) inflammation signaling of F4/80+ macrophages in iWAT. mRNA expression of genes responsible for (G) lipid metabolisms and (H) inflammatory cytokines and chemokines in HFD-fed WTBM→WT/HFD and KOBM→WT/HFD mice, *n*=6. Data were expressed as mean ± SEM, \**p* < 0.05.

### iRhom2 expression only in myeloid cells increases high fat diet-induced adipose inflammation and dyslipidemia

To better sustain the role of iRhom2 in MΦs in regulation of the pathogenesis of obesity, we next transplanted the bone marrow cells isolated from WT and/or iRhom2 KO mice to lethally irradiated iRhom2 KO mice to create a WTBM→KO mice. After treatment of HFD for 16 weeks, the WTBM→KO/HFD mice, in which iRhom2 was integrated only in myeloid cells, and KOBM→KO/HFD mice, in which iRhom2 was impaired in all cells, exhibited no significant difference in body weight and body composition (Fig S5 A and B). However, HFD-treated WTBM→KO/HFD mice showed increased severity in glucose tolerance and insulin resistance compared with those in HFD-fed KOBM→KO/HFD mice (Fig S5 C and D) as determined by GTT and ITT test, and AUC index. Furthermore, we subsequently investigated the alteration in insulin resistance associated indicators expression of iWAT in HFD-fed mice. Indeed, WTBM→KO/HFD mice displayed up regulated severity of insulin resistance signaling including p-IRS^Y608^, p-AKT, p-GSK3β, p-FOXO1 and PEPCK levels compared these in KOBM→KO/HFD mice (Fig S5 E). When status of inflammatory responses in iWAT were examined, significantly, WTBM→KO/HFD mice also exhibited increased TNF-α/NF-κB signaling activation, compared with corresponding mice, as indicated by western blotting analysis (Fig. S5 F). Meanwhile, determined by alterations in mRNA expression levels about lipid metabolism and adipose inflammation, WTBM→KO/HFD mice showed increased severity of dyslipidemia related indicators compared with KOBM→KO/HFD mice (Fig. S5 G). Consistently, WTBM→KO/HFD mice produced markedly more inflammation cytokines and chemokines in mRNA levels compared with those in KOBM→KO/HFD mice (Fig. S5 H). Taken together, the results presented in Fig. 4 and Fig. S5 significantly demonstrate a destructive role for iRhom2 in macrophages during HFD-triggered adipose inflammation and dyslipidemia.

## Discussion

The prevalence of obesity has increased dramatically in most industrialized countries over the past few decades. Obesity is a systemic disease that unleashes a prolonged low level of inflammation in insulin target organ including muscle, fat and liver, resulting in an overload and produce of a series of cytokines and chemokines that trigger systemic insulin resistance (28-31). Increasing studies forcefully indicate that obesity elevates the likelihood of numerous diseases, particularly cardiovascular and cerebrovascular diseases, type II diabetes mellitus, nonalcoholic fatty liver disease, obstructive sleep apnea, atherosclerosis, certain types of cancer, and osteoarthritis (32-36). Mechanistically, chronic inflammation infiltration serves as a critical role in obesity-stimulated metabolic disorder and various metabolism associated diseases (37). Unfortunately, the perspicuous molecular mechanisms by which how obesity lead to insulin resistance, systemic inflammation and metabolic diseases is still not fully understood. iRhom2, an inactive member of the rhomboid intramembrane proteinase family of serine proteases, has been treated as a key regulator in innate immunity (15-17, 38, 39). In the present study, we determined in a series of model animals’ experiments that adipose iRhom2 is involved in insulin resistance and inflammatory responses. By the different mice models of obesity, we further confirmed that iRhom2 plays a significantly accelerative role in obesity-induced sustained systemic metabolic disorder. Importantly, previous studies have suggested that iRhom2 is a crucial pathogenic mediator of inflammatory responses and required for the production of TNF-α secretion in immune cells (13, 16). The iRhom2 found in myeloid cells (MΦs) is required and essential for over-nutrition stress to stimulate or promote obesity and its associated metabolic disorder aspects in mice. In essence, over-activation by obesity dramatically elevates macrophages inflammatory levels, which in turn, up regulates adipocytes insulin resistance, further aggravates adipose inflammatory responses and chemotactic macrophages activation. Thus, we provided attractive demonstration to sustain that iRhom2 exhibits an adverse regulator in obesity by promoting insulin resistance and macrophage activation associated pro-inflammatory responses of fat tissues.

Our studies formerly illustrated that mice with sustained HFD feeding show obvious adipose inflammation and non-alcoholic fatty liver disease-like pathological phenotype compared with corresponding control mice (40, 41). In this current mice model of obesity, up regulation in the phosphorylation levels of NF-κB, TNF-α and TNFR1/2 in fat tissue, adipocytes, SVFs and macrophages were determined, demonstrating elevated activation events of downstream of iRhom2. Consistently, we also observed elevated phosphorylated NF-κB in *db/db* mice (a model of genetic obesity), which was accompanied by an up regulated adipose inflammation including over-activated downstream events of iRhom2. These results revealed a crosslink in iRhom2 signaling and obesity. Accordingly, in order to better sustain this, we confirmed the correlation mechanism of elevated iRhom2 activity to obesity associated metabolic disorder by investigation of indicators of metabolic syndrome. As well, we supposed a disadvantageous effect of iRhom2 in obesity-induced insulin resistance and adipose inflammation. On the other hand, we hoped that the loss of iRhom2 can alleviate the metabolic disorder caused by obesity. In this regard, indeed, iRhom2 deleted mice protected against HFD-induced obesity associated body over-weight, insulin resistance and abnormal lipid homeostasis, as well as increased adipocytes size and number compared with control groups. For adipose inflammation pathogenesis, macrophage infiltration significantly contributes to the development and process of metabolic dysfunction. Predictably, the iRhom2 deficiency restrained F4/80 positive macrophages activation in adipose tissues. Given that iRhom2 knockout was able to promote whole body glucose and lipid metabolism balance in HFD-induced obesity. We sought to confirm whether over-expression of iRhom2 accelerates obesity-triggered insulin resistance and dyslipidemia. iRhom2 over expression in mice were constructed using adenovirus associated virus 8 (AAV8) delivery system. In contrast to those in iRhom2 knockout mice, overactivation of iRhom2 results in the activity of its downstream TNFR1/2 signaling, which further phosphorylate and activate NF-κB, promoting expression of a series of inflammatory cytokines and chemokines in fat tissue, adipocytes and F4/80 positive macrophages such as TNF-α, IL-1β, IL-6, MCP-1, CCL1, IL-18, IL-17, IL-2, IL-10, IL-5, MIP-2, MIP-3β, MCP-3, PCT, HMGB1, MDC, GCP-2, CX3CL1, CXCL10 and CXCL12. Consistently, iRhom2 overactivation in HFD-fed mice markedly maintained lipid homeostasis disequilibria associated genes expression including increased cholesterol synthesis, fatty acid uptake and synthesis, and decreased fatty acid efflux and β-oxidation.

Previous studies have indicated that iRhom2 plays a key role in regulating the maturation of TNF-α convertase (ADAM17), which manages shedding of TNF-α by trafficking with ADAM17 and pro-inflammatory activity in vivo (12, 16). In view of such situation, to deeply highlight the essential role for iRhom2 in macrophage activation related insulin resistance and metabolic dysfunction, we afforded additional testimonies from BMT mice to sustain the harmful effect of iRhom2 in myeloid cells in progression and pathogenesis of obesity. In the functional deficit experiments, the BMT mice whose iRhom2 was deleted specifically in myeloid cells exhibited energetic glucose tolerance and reduced insulin resistance compared with mice whose iRhom2 was uninjured in all cells. Conversely, in functional acquisition experiment, BMT mice, which had uninjured iRhom2 specifically in myeloid cells, showed increased glucose intolerance and insulin resistance, further displayed elevated severity of HFD induced obesity aspects compared with mice whose iRhom2 was impaired in all cells. The comparative studies indicate that myeloid cells (macrophages)-specific iRhom2 is essential and required for the produce of cytokines in HFD-induced adipose inflammation and metabolic dysfunction. Of note, HFD-stimulated obesity phenotype in BMT mice whose iRhom2 was impaired only in myeloid cells showed a lower adipose inflammation and abnormal lipid metabolism related indicators levels than those in mice whose iRhom2 was intact in all cells, whereas HFD-stimulated obesity phenotype in BMT mice whose iRhom2 was not destructive only in myeloid cells exhibited a higher adipose inflammation and lipid metabolism disorder related indicators expression than those in mice whose iRhom2 was deleted in all cells. It should be pointed out that no significant differences of body weight and body composition in corresponding BMT animals participating in the comparative experiment. These data enrich a marked function from macrophage-specific iRhom2 in obesity associated adipose inflammation, lipid metabolism and insulin resistance than from iRhom2 served in other cells.

In conclusion, the current study identified a novel effect of iRhom2 on obesity and insulin resistance that is rely on the crosslink and interaction of iRhom2 with obesity induced macrophage activation, which enhances the TNFR1/2 and downstream events signaling activity of TNF-α and phosphorylated NF-κB cascades, accordingly leading to insulin resistance, adipose inflammation and dyslipidemia in response to an HFD challenge (Fig. 5). We further indicated that decrease or increase in macrophage-specific iRhom2 expression constitutionally displays a deleterious effect on adipose inflammation and insulin resistance and that this effect was confirmed by promoting macrophages pro-inflammatory activation. These conclusions strongly provide evidence sustaining the idea that iRhom2 may come up with a remarkable therapeutic target and a promising treatment approach for obesity and its associated metabolic diseases.

**Figure 5.**
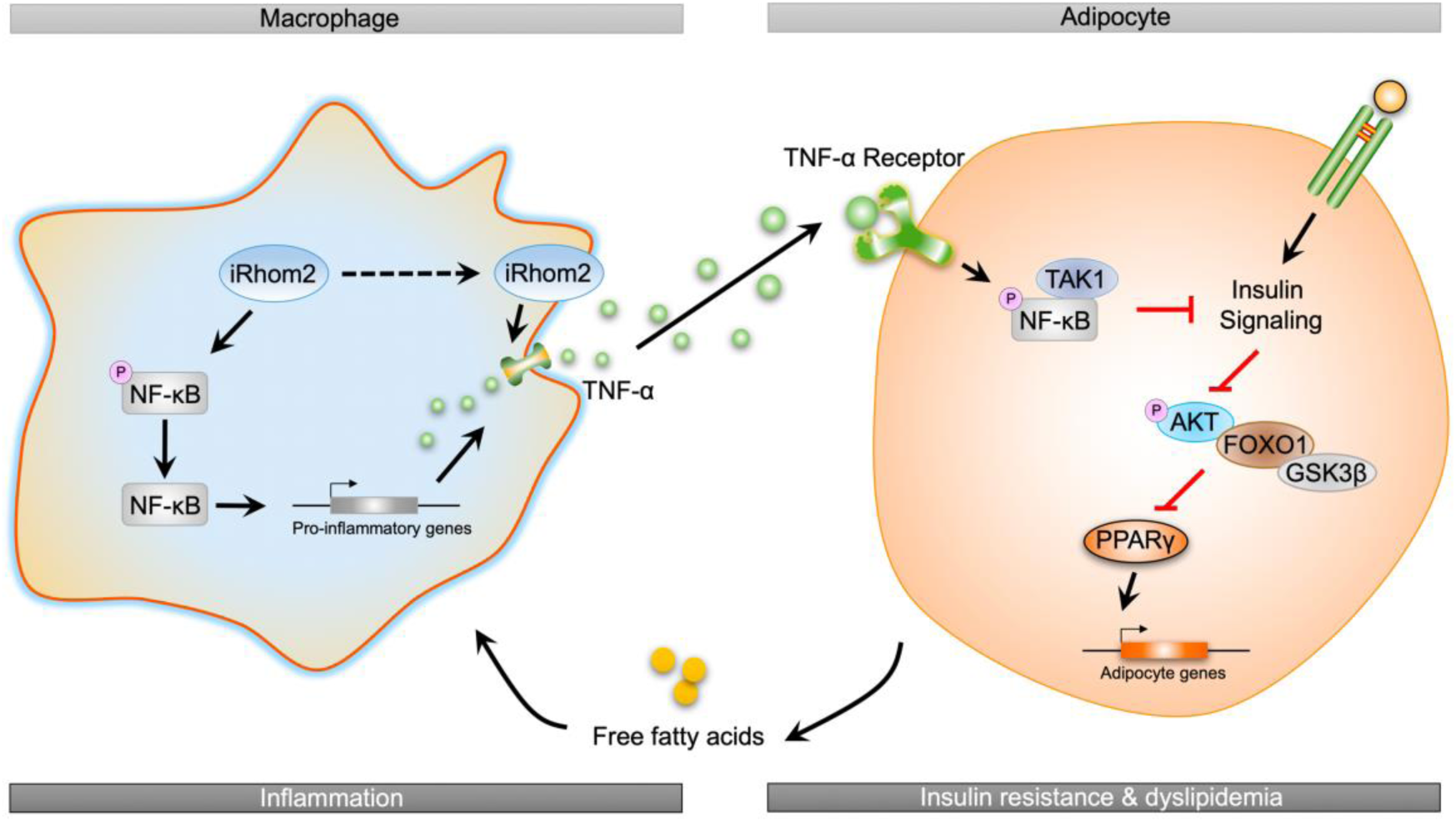
A possible schematic diagram displays the role of iRhom2 on high fat diet-induced adipose inflammation and insulin resistance.

## Materials and Methods

### Study Approval and Ethics Statement

All research procedures associating with mice were approved by the Institutional Animal Care and Use Committee in Chongqing Key Laboratory of Medicinal Resources in the Three Gorges Reservoir Region, School of Biological and Chemical Engineering, Chongqing University of Education, and were performed in accordance with the Guide for the Care and Use of Laboratory Animals, issued by the National Institutes of Health in 1996. The protocols used in this study were in accordance with the Regulations of Experimental Animal Administration issued by the Ministry of Science and Technology of the People’s Republic of China (http://www.most.gov.cn).

### Animal administration and experiment design

Inactive rhomboid-like protein 2 (iRhom2) deletion mice with C57BL/6N background were used in this work as described previously (17). Wild-type (WT) C57BL/6N mice were purchased from Beijing Vital River Laboratory Animal Technology Co., Ltd (Beijing, China). Prior to all experiments, the mice were allowed to adapt to the environment for 7 days. The mice were maintained in a constant temperature (controlled by GREE air conditioner), humidity and pathogen-free-controlled environment (25 °C ± 2 °C, 50% −60 %) with a standard 12 h light/12 h dark cycle, abundant food and water (pathogen-free) in their houses.

#### Model 1#

Male WT mice at the age of 6-8 weeks or *db/db* mice were accordingly constructed by feeding with high fat diet (HFD) fodder (20% kcal protein, 60 kcal% fat and 20% kcal carbohydrate, Cat#: D12492; Research Diets, New Brunswick, NJ, USA) for 16 weeks until experimental mice were sacrificed for further study. In addition, the mice that were fed a standard chow diet (20% kcal protein, 10 kcal% fat and 70% kcal carbohydrate, Cat#: D12450H; Research Diets, New Brunswick, NJ, USA) as control group (SCD). After the feeding period, mice were collected to investigate white adipose tissue signaling items downstream of iRhom2 associating with diet-induced adipose inflammation.

#### Model 2#

Male iRhom2 knockout (KO) mice and WT mice at the age of 6-8 weeks, were fed with HFD diet for 16 weeks to detect the alterations of iRhom2 expression, adipose inflammation and insulin resistance. Age-matched male KO mice and WT mice were separately fed with SCD fodder for 16 weeks and treated as controls.

#### Model 3#

Male WT mice at the age of 6-8 weeks were fed with HFD diet for first 4 weeks. Then, the adenovirus associated virus 8 (AAV8) delivery system was used to overexpress iRhom2 in the adipose tissue. Briefly, the opening reading frame (ORF) encoding iRhom2 without a stop codon was cloned into pAAV-CMV vector to generate pAAV-CMV-Rhbdf2-2A-EGFP-3FLAG-CW3SL. Accordingly, mice were injected with vector via tail vein with 100 μl of virus containing 2 × 10^11^ vg of AAV8 vectors and fed with HFD for additional 12 weeks. The empty vector based on pAAV-CMV was used to serve as control.

#### Model 4#

Male WT mice at the age of 6-8 weeks were lethally irradiated and then subjected to bone marrow transplantation (BMT) surgery with bone marrow cells isolated from iRhom2 KO mice and WT mice, and treated as WTBM→WT/HFD and KOBM→WT/HFD mice, respectively, as described previously (26, 27). After recovery for 4 weeks, these mice were fed with HFD for 16 weeks as described in model 2#.

#### Model 5#

Male iRhom2 KO mice at the age of 6-8 weeks were lethally irradiated and then subjected to bone marrow transplantation (BMT) surgery with bone marrow cells collected from iRhom2 KO mice and WT mice, and treated as KOBM→KO/HFD and WTBM→KO/HFD mice, respectively. After recovery for 4 weeks, these mice were fed with HFD for 16 weeks as described in model 2#.

During the feeding period, fasting blood glucose and insulin, body weight, food and water intake were recorded every 4 weeks or every week in model 2# mice. The body weight and body composition associating BMT mice were examined at first week and last week during experimental period. Serum of eye blood were harvested from all mice and stored at −20°C. Epididymal white adipose tissues (eWAT) and inguinal white adipose tissues (iWAT) were also carefully isolated and stored at −80°C or corresponding tissue analysis treatment fluid.

### Body composition, glucose and insulin tolerance detection

Mice body composition including lean mass, fat mass, relative percent of fat and relative bone mineral density were examined using Dual-energy X-ray absorptiometry (DEXA) (GE Medical Systems, China). For glucose tolerance test (GTT) and insulin tolerance tests (ITT) detection, all mice associating with this regard were fasted for 8 h to confirm the correction of physiological response. Mice were given an intraperitoneal injection of glucose (2 g/kg body weight) (Cat#: D810588, Macklin Inc., Shanghai, China). Then, the concentration of blood glucose of tail venous blood at 0 min, 15 min, 30 min, 60 min and 120 min after glucose treatment were examined using commercial blood glucose test strips (ACCU-CHEK®, Roche Diabetes Care GmbH, Shanghai, China). For insulin resistance detection, mice were treated with an intraperitoneal injection of insulin (1 U/kg body weight, Sigma Aldrich). The GTT and ITT analysis protocol were used in accordance with our previous study (41). Subsequently, blood samples were collected from the tail vein at 0 min, 15 min, 30 min, 60 min and 120 min post-injection for measurement of glucose levels. Homeostasis model assessment (HOMA)-IR was calculated from fasting levels of glucose and insulin in serum, respectively.

### Histological examination

The fat tissue samples from each group of mice were fixed with 4% paraformaldehyde, embedded in paraffin, and sectioned transversely. Adipose or liver tissues were subjected to hematoxylin and eosin (H&E) or oil red O staining to visualize the pattern of lipid accumulation and the impaired status of tissues. Briefly, the mice liver and white adipose tissues were fixed with 4% paraformaldehyde (Cat#: P1110, Solarbio Life Sciences, Beijing, China), embedded in paraffin (Cat#: P100928, Aladdin, Shanghai, China), and sectioned transversely. Thin sections were stained with H&E (Hematoxylin and Eosin Staining Kit, Yeasen, Shanghai, China) according to a standard histopathological processes. All sections were detected by 3 histologists without knowledge of the treatment procedure. To visualize lipid accumulation, tissues were frozen in Tissue-Tek OCT (Tissue-Tek, Sakura Finetek, USA) and sections were then stained with Oil Red O Stain Kit (Cat#: G1260, Solarbio Life Sciences, Beijing, China) for 10 min. After being rinsed with 60% isopropyl alcohol (Cat#: I811925, AR, Macklin Inc., Shanghai, China), the tissue sections were re-stained with haematoxylin. For immunohistochemistry assay, embedded sections were deparaffinized before administration with primary antibodies at 4 °C overnight. The tissues were subjected to immunohistochemical (IHC) staining for the measurement of F4/80 (Abcam, ab100790, dilution 1:300) expression. All the histological procedure was performed in accordance with the standard procedures as indicated in reagent specifications. All the images were captured using optical microscope (Olympus, Japan).

### Adipocytes size and number detection

Adipose cells and number detection were performed according to the protocol described previously (42). The average of cells diameter was examined using Image J software derived from National Institutes of Health (NIH), USA. Total adipose cells numbers were determined by counting cells on H&E sections from at least 3 fields per mouse. The size of cells was approximated assuming cubic packing as indicated previously (42).

### F4/80 positive macrophages collection in adipose tissues

The protocol of F4/80 positive (F4/80+) macrophages collection in adipose tissues was performed according to the previous studies with a certain modification (7, 43) and manufacturer’s instructions. Isolation and purification of F4/80+ macrophages (MΦ) was collected from stromal vascular fraction (SVF) of adipose tissues based on PE magnetic particles separation as indicated in the manufacturer’s specification. In brief, 2 × 10^7^ SVF cells was re-suspended in 120 μl Hanks buffer (Solarbio, Beijing, China) and then stained with 15 μl mouse PE-conjugated F4/80 mAbs (Cat#: FAB5580P-100UG, R&D system), and co-incubated with 30 μl anti-PE magnetic beads for 20 min. Next, the washed cells were re-suspended in 1 ml Hanks buffer and then subjected to XS Separation columns (Miltenyl Biotec MACS). The positive MΦ cells were eluted and harvested for further experiments.

### Real-time quantitative PCR and immunoblotting assay

TRIzol production (Cat#:15596026, Invitrogen, Thermo Fisher Scientific, USA) was used to collect the total RNA extraction from cells or adipose tissues. In brief, 1 μg of total RNA samples were reverse transcribed using the M-MLV-RT system (Invitrogen™, Shanghai, China). The program was performed at 42 °C for 1 h and terminated by deactivation of the enzyme at 70 °C for 10 min. PCR were conducted using SYBR Green (Bio-Rad) in the ABI PRISM 7900HT detection systems (Applied Biosystems, USA). Pre-denatured products for amplification was performed at 94 °C for 55s; followed by 45 cycles at 95 °C for 30s, 57 °C for 30s and 73 °C for 30s; followed by 95 °C for 10s, 65 °C for 45s, and 40 °C for 60s. Fold induction values were calculated according the 2^-ΔΔCt^ expression, where ^Δ^Ct represents the differences in cycle threshold number between the target gene and GAPDH, and ^ΔΔ^Ct represents the relative change in the differences between control and treatment groups. The primer sequences for RT-PCR were produced by Sangon Biotech (Shanghai) Co., Ltd., and indicated in Table S1.

For western blotting examination, cells or tissues were homogenized into 10% (wt/vol) RIPA lysis buffer (pH 8.0, 1mM EDTA, 50 μg/ml aprotinin, 25mM Tris-HCl, 5 μg/ml leupeptin, 1mM Pefabloc SC, 4mM benzamidine, 5 μg/ml soybean trypsin inhibitor) to yield a homogenate. Next, the final liquid supernatants were harvested by centrifugation at 13500 rpm, 4 °C for 30 min. Protein concentration was measured using Pierce™ Rapid Gold BCA Protein Assay Kit (Thermo, USA) with bovine serum albumin as a standard. The total protein extraction will be performed for western blot analysis. Equal amounts of total protein of tissues were subjected to 10%-12% SDS-PAGE and then transferred to a 0.45 μM PVDF membrane (Millipore Company, USA) followed by immunoblotting using the primary antibodies. The primary antibodies used in this study including anti-GAPDH (Abcam, ab8245, dilution 1:5000), anti-Tubulin (Abcam, ab7291, dilution 1:5000), anti-p-IRS-1(Y608) (Millipore, dilution 1:1000), IRS-1 (Abcam, ab52167, dilution 1:1000), p-AKT (CST, 4060, dilution 1:1000), AKT (CST, 4691, dilution 1:1000), p-GSK3β (CST, 9322, dilution 1:1000), GSK3β (CST, 12456, dilution 1:1000), FOXO1 (CST, 2880, dilution 1:1000), p-FOXO1 (Abcam, ab131339, dilution 1:1000), PEPCK (Santa-Cruz, sc-271029, dilution 1:1000), anti-NF-κB (Abcam, ab16502, dilution 1:1000), anti-p-NF-κB (Abcam, ab86299, dilution 1:1000), anti-Rhbdf2 (Biobyt, orb386934, dilution 1:1000), anti-TNF-α (CST, 3707, dilution 1:1000), anti-TNFR1 (Abcam, ab19139, dilution 1:1000), anti-TNFR2 (Abcam, ab109322, dilution 1:1000), anti-IκBα (Abcam, ab32518, dilution 1:1000), anti-p-IκBα (CST, 9246, dilution 1:1000), anti-TAK1 (Abcam, ab109526, dilution 1:1000) and anti-p-TAK1 (CST, 9339, dilution 1:1000). Accordingly, the membranes were blocked with 5% skim milk (DifcoTM Skim Milk, BD, USA) in 1×TBS buffer (Cat#: T1080, Solarbio, Beijing, China) containing 0.1% Tween-20 (1247ML100, BioFROXX, Germany) (TBST) for 1 h and incubated with the primary antibodies in fridge overnight (4°C). Membranes were rinsed in 1×TBST and incubated with HRP-conjugated anti-rabbit or anti-mouse secondary antibodies (Abcam) for 1-2 h at room temperature (25°C). Bands were visualized by Enhanced New Super ECL Western Blotting Substrate (KeyGEN BioTECH, China) and exposed to Kodak (Eastman Kodak Company, USA) X-ray film. Corresponding protein expression will be determined as grey value (Version 1.52g, Mac OS X, Image J, National Institutes of Health, USA) and standardized to housekeeping genes (GAPDH) and expressed as a fold of control.

### Measurement of cytokines and chemokines produce

The cytokines and chemokines of mice were detected using corresponding commercial ELISA kit. The TNF-α (Cat#: MTA00B), IL-1β (Cat#: MLB00C), IL-6 (Cat#: M6000B), IL-18 (Cat#: DY122-05), MCP-1 (Cat#: SMJE00B), CCL1 (Cat#: DY845), IL-17 (Cat#: SM1700), IL-2 (Cat#: SM2000), IL-10 (Cat#: SM1000B), IL-5 (Cat#: SM5000), MIP-2 (Cat#: SMM200), MIP-3β (Cat#: DY440), MDC (Cat#: MCC220), CX3CL1 (Cat#: MCX310), CXCL10 (Cat#: DY466-05) and CXCL12 (Cat#: DY460) ELISA kits were purchased from R&D system (Shanghai, China) and used according to the product specification. The MCP-3 (Cat#: BMS6006INST) ELISA kit were from Invitrogen. The GCP-2 (Cat#: ab100719) kit was obtained from Abcam. The HMGB1 ELISA kit (Cat#: 326054329) was purchased from Shino-Test Corporation, Kanagawa, Japan. The procalcitonin ELISA kit (Cat#: EF014005) was from Life Science. All the corresponding serum were carefully stored at −80°C refrigerator until used.

### Statistical analysis

Data were expressed as mean ± standard error of the mean (SEM). Comparisons between every groups were analyzed by one-way ANOVA followed by Dunnett’s multiple comparisons test or unpaired 2-tailed Student’s *t* test. GraphPad Prism Software (version 7.0 for Mac OS X Snow Leopard; Graph Pad Software, Inc., San Diego, CA) was used for the analysis. A p-value less than 0.05 will be considered significant.

## Acknowledgments

This work was supported by National Natural Science Foundation of China (NSFC Grant number: 81703527); Chongqing Research Program of Basic Research and Frontier Technology (Grant number: cstc2017jcyjAX0356, cstc2018jcyjA3686, cstc2018jcyjA1472 and cstc2018jcyjA3533); 2018 Chongqing College Students’ Innovation and Entrepreneurship Training Project (Grant number: 201814388021 and 201814388022); School-level Research Program of Chongqing University of Education (Grant number: KY201710B and 17GZKP01) and Advanced Programs of Post-doctor of Chongqing (Grant number: 2017LY39). We also thank LetPub (www.letpub.com) for its linguistic assistance during the preparation of this manuscript.

